# The COVID-19 RNA vaccine Spikevax^TM^ transfects human umbilical endothelial cells and induces Spike protein production, inflammation and leukocyte adhesion

**DOI:** 10.64898/2026.09.15.751712

**Authors:** Marco Cosentino, Marco Ferrari, Nicola Schiavone, Emanuela Rasini, Alessandra Luini, Massimiliano Legnaro, Maurizio Federico, Franca Marino, Giovanni Vanni Frajese

## Abstract

Endothelial cell dysfunction plays a key role in the pathogenesis of severe and critical COVID-19, leading to a multi-systemic inflammatory disease, and results from the direct interaction of the SARS-CoV-2 Spike proteins with endothelial cells. COVID-19 RNA vaccines encode a recombinant SARS-CoV-2 Spike protein which undergoes systemic biodistribution. No information is however so far available on the direct effects of COVID-19 RNA vaccines on human endothelial cells. In the present study, we exposed cultured human umbilical venous endothelial cells (HUVEC) to the COVID-19 RNA vaccine Spikevax^TM^ (Moderna, Inc.), and thereafter we measured the expression of the Spike protein, of the proinflammatory cytokines interleukin (IL)-6 and tumor necrosis factor (TNF)-α, of the adhesion molecules intercellular adhesion molecule 1 (ICAM-1) and vascular cell adhesion molecule 1 (VCAM-1), as well as the attachment of human leukocytes to HUVEC layers. We also assessed the effects of Spikevax^TM^ on HUVEC viability. Exposure of HUVEC to the COVID-19 RNA vaccine Spikevax^TM^ resulted in effective cell transfection, and subsequent production of the Spike protein, which was expressed in the cells and secreted in the culture medium. Spike protein production was accompanied by increased gene expression of IL-6 and TNF-α and of ICAM-1 and VCAM-1, as well as by increased attachment of leukocytes to HUVEC monolayers. Exposure to the COVID-19 RNA vaccine Spikevax^TM^ did not affect HUVEC viability. Our results provide a mechanistic explanation to post-COVID-19 RNA vaccination pathologies resulting from endothelial dysfunction, such as during atherosclerosis, autoimmune inflammation, and systemic inflammatory conditions.

## Introduction

Coronavirus disease 2019 (COVID-19) is caused by severe acute respiratory syndrome coronavirus 2 (SARS-CoV-2). The natural course of SARS-CoV-2 infection is characterized by a viral pneumonia presenting as fever and progressively worsening cough; however, some patients may subsequently develop acute respiratory distress syndrome (ARDS) requiring intensive care (Zhu et al., 2020; Huang et al., 2019; Berlin et al., 2020). Death from severe COVID-19 is usually due to pulmonary edema, cytokine storm, multiple organ failure, and disseminated intravascular coagulation (Elezkurtaj et al., 2021). Widespread endothelial cell dysfunction plays a key role in the pathogenesis of severe and critical COVID-19, leading to a multi-systemic inflammatory disease, characterized by thrombo-embolic phenomena, and microcirculatory dysfunction (Otifi et al., 2022; Bonaventura et al., 2021; Nägele et al., 2020).

Endothelial cell dysfunction in COVID-19 has been initially regarded as a consequence of the strong inflammatory response triggered by SARS-CoV-2. Increasing evidence however indicates that direct interaction of SARS-CoV-2 with endothelial cells through the Spike protein is pivotal in the pathogenesis of endothelial dysfunction occurring in COVID-19 (Perico et al., 2024). The SARS-CoV-2 Spike protein directly binds the angiotensin-converting enzyme 2 (ACE2) receptor on human endothelial cells, resulting in several detrimental effects, including: ACE2 downregulation and mitochondrial dysfunction (Varga et al., 2020), increased production of interleukin (IL)-6, MCP-1, intercellular adhesion molecule 1 (ICAM-1) and PAI-1, and NFkB activation (Montezano et al., 2023), increased intracellular reactive oxygen species (ROS) levels, which inhibit the PI3K/AKT/mTOR pathway and induce autophagy and apoptosis (Li et al., 2021), induced endothelial cell permeability and von Willebrand factor secretion (Guo and Kanamarlapudi, 2023), increased ROS generation, vascular cell adhesion molecule 1 (VCAM-1), ICAM-1, and leukocyte attachment (Meyer et al., 2021), activated NF-κB, promoted pro-inflammatory cytokines release, triggered the priming and activation of the NLRP3 inflammasome system, and enhanced production of coagulation factors such as von Willebrand factor, factor VIII or tissue factor (Villacampa et al., 2024), increased expression of ICAM-1 and E-selectin, release of cytokines and chemokines, and in particular of IL-6, CXCL1 and CXCL2, increased leukocyte adhesion and clot formation (Gultom et al., 2024). Additional targets of the Spike protein on human endothelial cells include the integrin *a*5β1, resulting in induced nuclear translocation of NF-κB, subsequent expression of VCAM1 and ICAM1, coagulation factors, proinflammatory cytokines such as tumor necrosis factor (TNF)-α, IL-1β, and IL-6, increased adhesion of peripheral blood leukocytes and increased permeability (Robles et al., 2022).

Mass vaccination against SARS-CoV-2 has been the main strategy implemented worldwide to mitigate the COVID-19 pandemic. In most Western countries, COVID-19 vaccination campaigns, were based on RNA vaccines encoding a recombinant SARS-CoV-2 Spike protein with specific amino acid variations introduced to maintain the protein in a prefusion state and uncleavable form (Teo, 2022). COVID-19 RNA vaccines were presented as intrinsically safe, based on the idea that, similar to conventional vaccines, after intramuscular injection, most of the dose would remain in the muscle and the rest would drain through the lymphatic system, being eventually captured by antigen-presenting cells and B cells and undergoing complete elimination in a few dozen hours at most (Lindsay et al., 2019; Lowe, 2021). However, convincing evidence has rapidly accumulated supporting extensive systemic disposition of both vaccine-derived SARS-CoV-2 S protein RNA and the resulting Spike protein (reviewed in: Trougakos et al., 2022; Cosentino and Marino, 2022), implying that vaccine-derived RNA translation may occur in principle in any tissues and organs, leading to the production of unpredictable amounts of Spike protein. On this basis, many authors have proposed that inappropriate production of Spike protein in vulnerable tissues may represent a major risk factor for local tissue damage, leading to adverse effects depending on the location and amount of Spike protein expression (or of local distribution from general circulation) (Cosentino and Marino, 2022; Trougakos et al., 2022; Parry et al., 2023). This hypothesis, however, is based almost entirely on what is known about the toxicity of the viral Spike protein, since few studies have so far directly examined the effects of COVID-19 RNA vaccines on human cells and tissues.

To obtain direct evidence about the effects of COVID-19 RNA vaccines on human endothelial cells, we exposed cultured human umbilical venous endothelial cells (HUVEC) to the COVID-19 RNA vaccine Spikevax^TM^, manufactured by Moderna Inc. (European Medicines Agency, 2021), and thereafter we measured the expression of the Spike protein, of proinflammatory cytokines such as IL-6 and TNF-α, and of the adhesion molecules ICAM-1 and VCAM-1, as well as the attachment of human leukocytes to HUVEC layers. We also assessed the effects of Spikevax^TM^ on HUVEC viability.

## Materials and methods

### Antibodies and reagents

Ficoll-Paque Plus was purchased from GEhealthcare (GeHealthcare, Uppsala, Sweeden). Lipopolysaccharide (LPS) from *Escherichia coli* (LPS) was obtained from Sigma-Aldrich (cod. L3137-5 mg, Sigma-Aldrich Srl, MI, Italy). Recombinant human CXCL8/IL-8 were purchased from R&D System (R&D System, MN, USA). APC-H7 – conjugated mouse anti-human CD45 (cod. 560178, clone 2D1, mouse IgG1,k) was obtained from Becton Dickinson, (BD Italy S.p.A., Milan). Human umbilical vein endothelial cells (HUVEC) and Endothelial Cell Growth Kit were purchased from PromoCell (PromoCell Gmbh, Germany), and the EndoGRO™ VEGF Complete Media Kit from Millipore (Millipore S.p.A., MI, Italy). The COVID-19 RNA vaccine Spikevax^TM^ (Moderna Inc., Cambridge, MA, U.S.A., batch number 000030A) was a kind gift from the National Center for Global Health, Istituto Superiore di Sanità, Viale Regina Elena, 299, Rome (Italy).

### HUVEC culture

Experiments were performed on human umbilical vein endothelial cells (HUVEC). HUVEC were purchased from PromoCell (cod. C-12205, PromoCell GmbH) and routinely cultured in endothelial cell growth medium kit (cod. C-22110, PromoCell GmbH) containing fetal calf serum (FCS) (0,02 ml/ml), endothelial cell growth supplement/heparin (ECGS/H) (0,004 ml/ml), human epithelial growth factor (hEGF) (0,1 ng/ml), human basic fibroblast growth factor (hb-FGF) (1 ng/ml), and hydrocortisone (1 µg/ml) (Endothelial Cell Growth Complete Medium). The culture flasks were maintained at 37°C in a humidified atmosphere of 5% CO2. HUVEC used for the experiments were between passage 8 and 15.

### HUVEC viability

Cell viability assay was performed by flow cytometry and annexin V (ANX)/propidium iodide (PI) staining, as previously described (Fanelli et al, 2014). FACSCelesta™ flow cytometer (Becton Dickinson Italy, Milan, Italy) and data were analyzed using BD FACSDiva software (version 8.0.1.1). HUVEC were identified on the basis of forward-scatter (FSC) and side-scatter (SSC) properties, and a minimum of 20000 cells for each sample was collected in the gate. Viable, apoptotic and necrotic HUVEC were identified on a biparametric plot ANX-FITC vs PI. Data were finally expressed as% viable (ANX-/PI-), early apoptotic cells (ANX+/PI-), late apoptotic/necrotic cells (ANX+/PI+) and necrotic cells (ANX-/PI+). The experimental designs used to assess HUVEC viability is outlined in **Supplementary Figure 1**.

### RNA isolation and real-time PCR assays

Total RNA was extracted from 5 x 10^4^ HUVEC using a GeneJET RNA Purification Kit (Thermo Fisher Scientific, Waltham, MA, USA). The amount of extracted RNA was estimated by spectrophotometry at λ = 260 nm and reverse-transcribed using a random primer and a High-Capacity cDNA RT Kit (Thermo Fisher Scientific). cDNA amplification was performed using TaqMan™ Universal PCR Master Mix (Thermo Fisher Scientific) with specific primers and probes for the RNA sequence encoding the COVID-19 RNA vaccine Spikevax^TM^-induced Spike protein, and for IL-6, TNF-α, ICAM1 and VCAM1 (Thermo Fisher Scientific), and assayed on a StepOne® System (Thermo Fisher Scientific, Waltham, MA, USA). For further details, see **Supplementary Table 1.** Assays were performed in triplicate for each sample, and mRNA levels were expressed as 2^−ΔCt^, where ΔCt = [Ct (sample) − Ct (housekeeping gene)]. Relative expression was determined by normalizing towards the expression of *RPS18*, which encodes for 18S cDNA. Data analysis was performed using StepOne Software™ 2.2.2 (Thermo Fisher Scientific). The experimental designs used to investigate the mRNA levels of the Spike protein is shown in **Supplementary Figure 2**, and of proinflammatory cytokines and adhesion molecules is depicted in **Supplementary Figure 3**.

### ELISA assay of SARS-CoV-2 Spike protein

Spike protein levels were measured in HUVEC supernatants using a commercial enzyme-linked immunosorbent assay (ELISA) kit (Invitrogen, code: EH492RB), according to the protocol supplied by the manufacturer. The experimental designs used to assess Spike protein levels in HUVEC supernatants is described in **Supplementary Figure 2**.

### Western Blot assay of SARS-CoV-2 S protein

Aliquots of 40 μg of HUVECs whole-cell lysates in RIPA buffer with protease inhibitors (cOmplete ULTRA Tablets, Roche, Basel, Switzerland, code: 05892791001), were subjected to Western blotting. Protein extracts were separated by SDS-PAGE (Thermo Fisher Scientific) and transferred onto nitrocellulose membranes (Bio-Rad, Hercules, CA, USA). Membranes were incubated in Odyssey Blocking Solution (Merck Millipore, Milan, Italy) for 1 h at room temperature. Membranes were then incubated overnight at 4 °C with the primary antibody (all primary antibodies were used diluted 1:1000 in a mix of 1:1 Odyssey Blocking Solution and PBS-Tween 0.1%), washed with PBS-Tween 0.1% solution, and probed with the secondary IRDye antibody according to the manufacturer’s instructions (secondary antibody diluted 1:12,000). Primary and secondary antibodies: anti-Sars-COV-2 Spike/S2 (40590-T62, Sino Biological, Beijing, China), anti-α-actin (MA1-744, Invitrogen, Carlsbad, CA, USA), Goat anti-rabbit IRDye 800, Goat anti-mouse IRDye 800 (926-32211 and 926-32210, LI-COR Biosciences, Lincoln, NE, USA). The protein bands were analyzed by the Odyssey Infrared Imaging System (LI-COR Bioscience).

### Leukocyte to HUVEC adhesion assay

The adhesion of leukocytes to HUVEC monolayers under static conditions was quantified by a flow cytometry assay, according to a previously published procedure (Marino et al., 2017). Briefly, HUVEC, maintained as above described, were plated in a 12-well plate at a density of 0.2 × 10^6^/ml in the medium. After 24 h cells reached 80–90% confluency and formed a monolayer that was left untreated (medium alone) or treated with 1 µg/ml of the COVID-19 RNA vaccine Spikevax^TM^ for 48 h. In each experiment, a HUVEC monolayer stimulated with 1 µg/ml LPS for 24 h was included as positive control for leukocyte adhesion. HUVEC monolayers were carefully washed two times with sterile PBS and 1 × 10^6^ CD45-labeled leukocytes (1 ml of cell suspension) were added. Samples were incubated for 30 min at 37 °C. After, incubation the medium was removed to eliminate nonadherent leukocytes, and HUVEC monolayers were carefully washed three times with sterile PBS. Finally, HUVEC with adhering leukocytes were detached by using a non-enzymatic solution, cell suspensions were centrifuged, and supernatants were removed. The pellets were resuspended in 350 μl PBS and kept on ice until flow cytometric acquisition. For the evaluation of leukocyte adhesion to HUVEC the same gating strategy described in Marino et al. (2017) was applied.

### DNA isolation from HUVEC and search for the Spike protein gene sequence

DNA was obtained from HUVEC using a standard DNA extraction protocol (Thermo Fisher Scientific, code: 4403319). Briefly, 2 µl of the cell culture suspension was incubated for 3 minutes with 20 µl of lysis solution at room temperature; then, 20 µl of DNA stabilizing solution was added. To eliminate RNA contamination, genetic material was incubated for 30 min with 10 µl Genomic RNase A (Thermo Fisher Scientific, code: R1253). Samples were stored at −20°C until real-time PCR analysis. Real-time PCR was performed using 10 ng of DNA from HUVEC cells incubated for 2, 5, 24, 48, 72 or 144 h with 1 µg/mL of the COVID-19 RNA vaccine Spikevax^TM^. The procedure included the following steps: 30 min at 50°C for reverse transcription, followed by 40 cycles of 15 s at 95°C and 1 min at 60°C, and a final 2-min extension period at 95°C. Previously validated primer pairs for Spikevax^TM^ (Hanna et al., 2022), and for the housekeeping gene Glyceraldehyde-3-phosphate dehydrogenase (GAPDH) (Aldén et al., 2022) were used with Maxima SYBR Green/ROX qPCR Master Mix (Thermo Fisher Scientific, Waltham, MA, USA). The positive control consisted of 5 µg/ml of cDNA obtained from the reverse transcription of the COVID-19 RNA vaccine Spikevax^TM^, added to HUVEC at the end of the incubation period. The negative control consisted of HUVEC alone.

## Results

### Spike protein expression in HUVEC cultured in the presence of the COVID-19 RNA vaccine Spikevax^TM^

Incubation of HUVEC for up to 144 h with the COVID-19 RNA vaccine Spikevax^TM^ either at the concentration of 1 or 3 µg/ml did not affect cell viability. Flow cytometry analysis showed that either concentrations did not affect the percentage of viable, early apoptotic, late apoptotic or necrotic cells in comparison to control conditions (**Supplementary Table 2**).

Exposure of HUVEC to 1 or 3 µg/ml of RNA of the COVID-19 RNA vaccine Spikevax^TM^ induced a time- and concentration-dependent production of the Spike protein, which was identified in the cells as both RNA and protein, and in the supernatant as protein (**Figure 1**, **Figure 2** and **Supplementary Figure 5**). In particular, the full-length Spike protein is visible as a band below 260 kDa, while a smaller-size band of approximately 130 kDa may represent the S2 cleavage product (**Figure 2** and **Supplementary Figure 5**).

**Figure 1.**
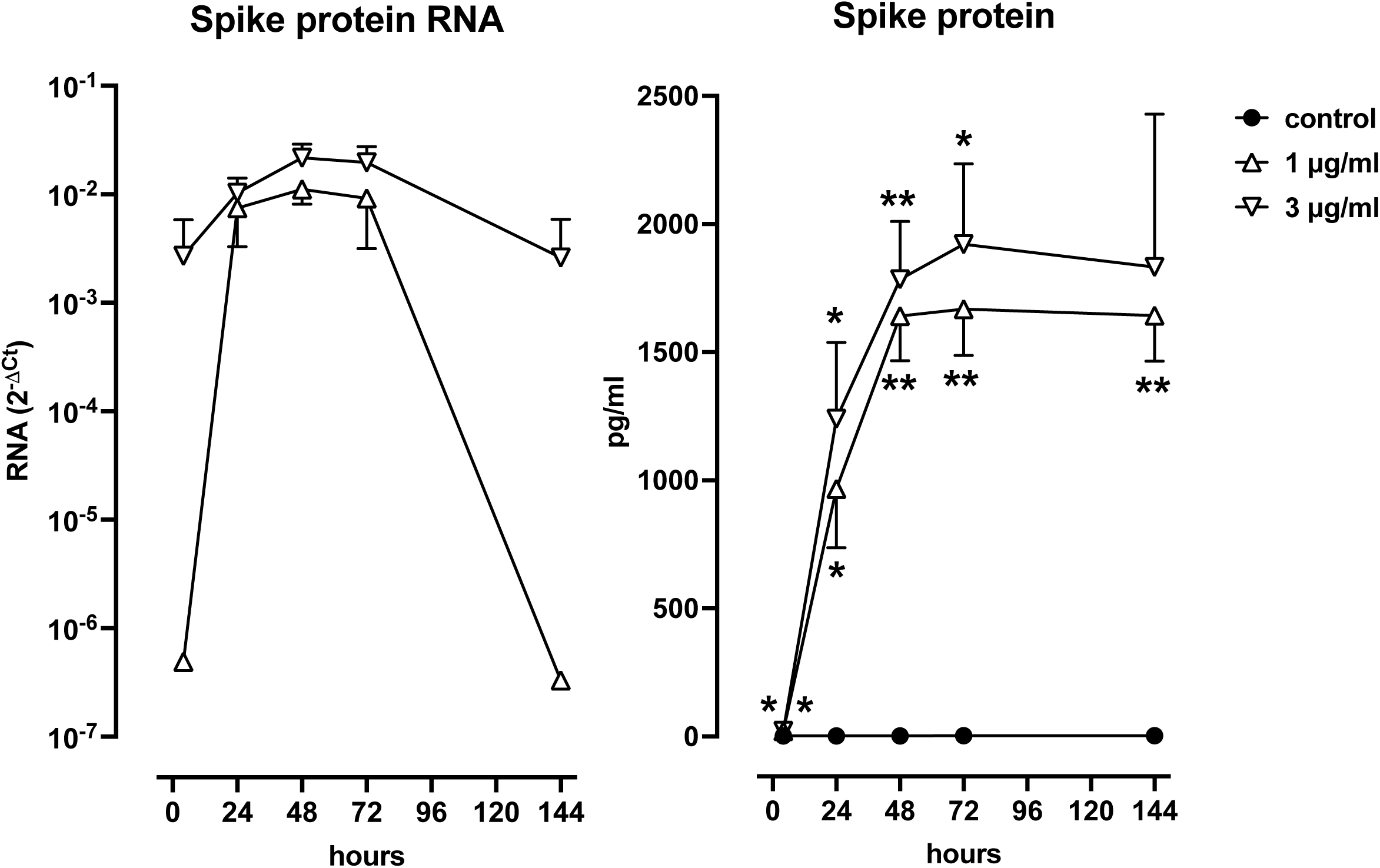
Spike protein expression in HUVEC cultured in the presence of the COVID-19 RNA vaccine Spikevax^TM^ at different concentrations. Left: RNA levels assayed by real time PCR. Right: protein levels measured by ELISA. Data are shown as mean±SD of 3-4 experiments. * = P<0,05 and ** = P<0,01 vs control. Left graph does not include control as RNA levels in the absence of Spikevax^TM^ were always undetectable.

**Figure 2.**
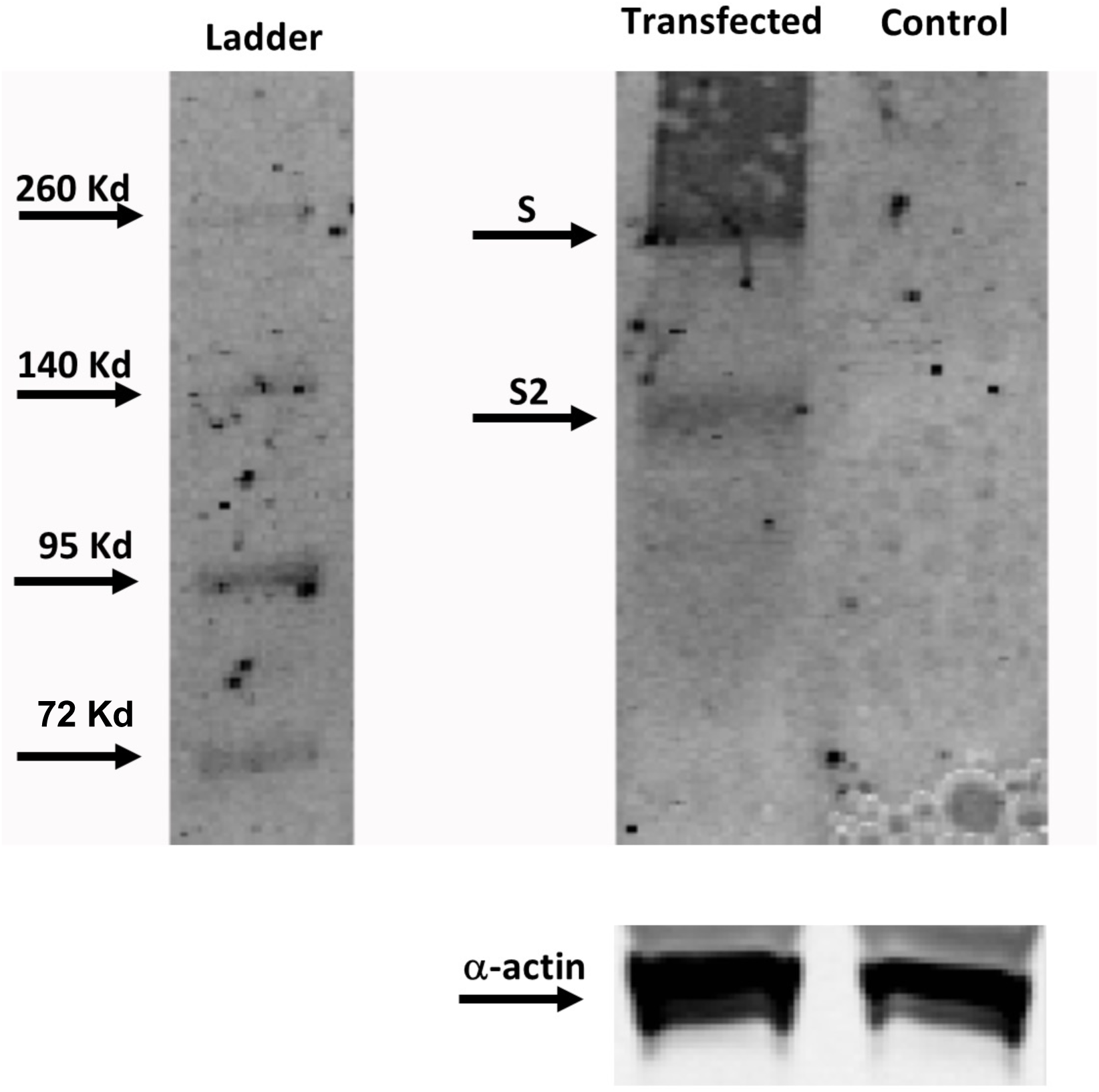
Western blot analysis of 40 μg of total protein extract from HUVEC treated for 48 h with the COVID-19 RNA vaccine Spikevax^TM^ at the concentration of 3 µg/ml. Control: untreated HUVEC. S: full-length Spike protein. S2: Spike protein S2 subunit.

Transfection was already evident after 4 h exposure with both 1 and 3 µg/ml, however Spike protein RNA levels were about 10.000 times higher with 3 µg/ml. In the presence of 3 µg/ml, Spike protein RNA levels remained high even at 144 h, when those induced by 1 µg/ml began to decline (**Figure 1, left panel**).

The Spike protein was detectable in the supernatant in trace amounts at 4 h with 1 or 3 µg/ml of COVID-19 RNA vaccine, and increased thereafter, peaking at 72 h and subsequently remaining constantly high. Spike protein concentrations from 72 h onwards were on average 1.643-1.668 pg/mL with 1 µg/ml, and 1.832-1.921 pg/mL with 3 µg/ml (**Figure 1, right panel**).

### Increased mRNA levels for proinflammatory cytokines and adhesion molecules in HUVEC cultured in the presence of the COVID-19 RNA vaccine Spikevax^TM^

Exposure of HUVEC to 1 or 3 µg/ml of the COVID-19 RNA vaccine Spikevax^TM^ induced a significant increase of the mRNA levels for the proinflammatory cytokines IL-6 and TNF-α as well as for the adhesion molecules ICAM-1 and VCAM-1 (**Figure 3**). Adhesion molecules mRNA significantly increased in the first 24 h, while TNF-α mRNA significantly increased up to 48 h and IL-6 mRNA significantly increased up to 72-144 h (**Figure 3**).

**Figure 3.**
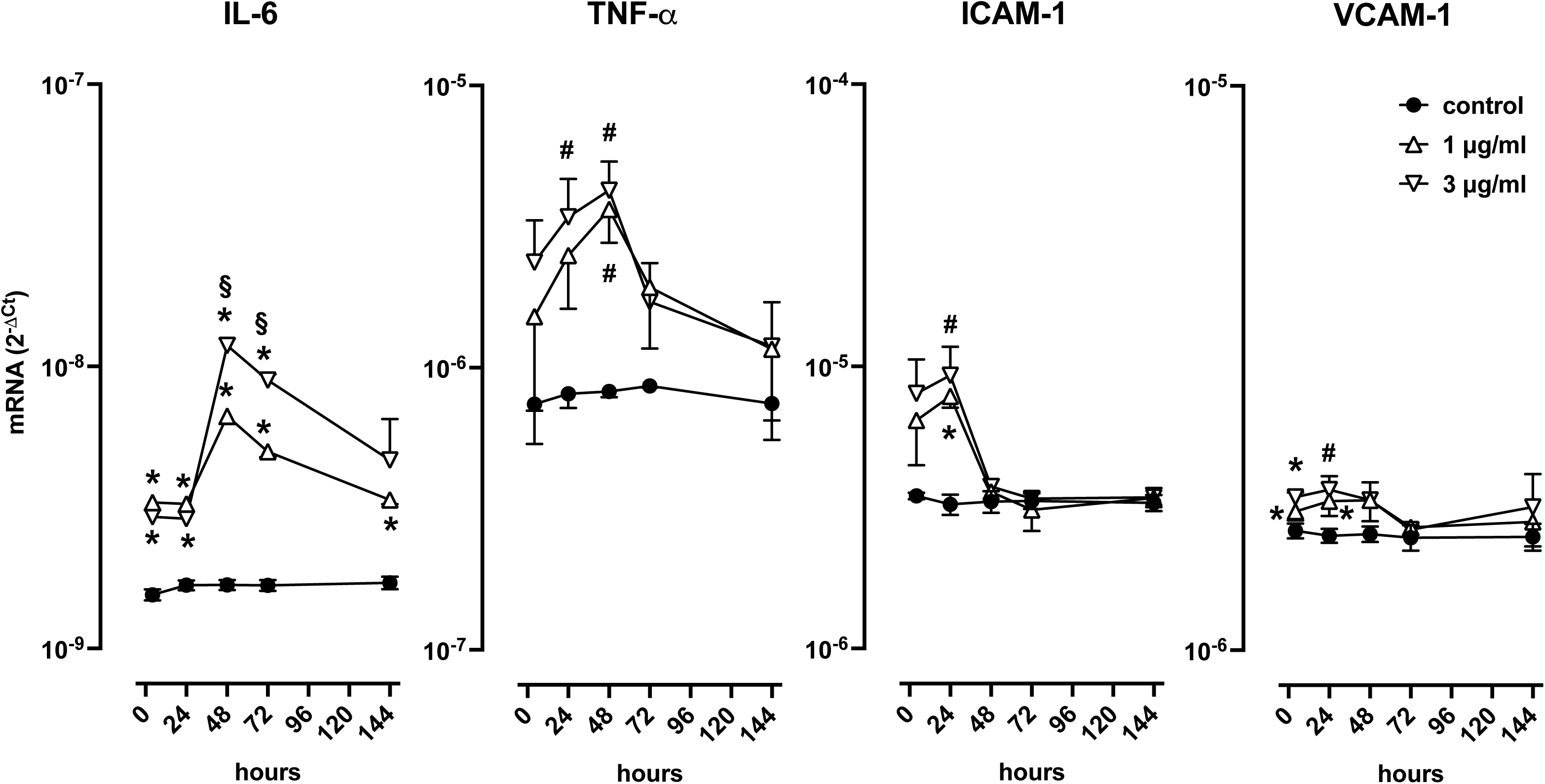
Effect of the COVID-19 RNA vaccine Spikevax^TM^ on the mRNA levels of the proinflammatory cytokines IL-6 and TNF-*a* and of the adhesion molecules ICAM-1 and VCAM-1 in HUVEC monolayers. Data are shown as means±SD of 4 experiments. # = P<0,05 and * = P<0,01 vs control and § = P<0,01 vs 1 µg/ml.

### Leukocyte adhesion to HUVEC cultured in the presence of the COVID-19 RNA vaccine SpikevaxTM

HUVEC were pretreated for 24 h with the COVID-19 RNA vaccine Spikevax^TM^ at the concentration of 1 µg/ml, and thereafter peripheral blood leukocytes were added to HUVEC and cocultured for an additional 48 h. In control experiments, no vaccine pretreatment was performed and LPS 1 µg/ml was added as positive control. Elsomeran significantly increased both monocyte and granulocyte adhesion to HUVEC respectively on average by about 29% and 47%. For comparison, LPS increased monocyte and granulocyte adhesion by 163% and 167%, respectively (**Figure 4**).

**Figure 4.**
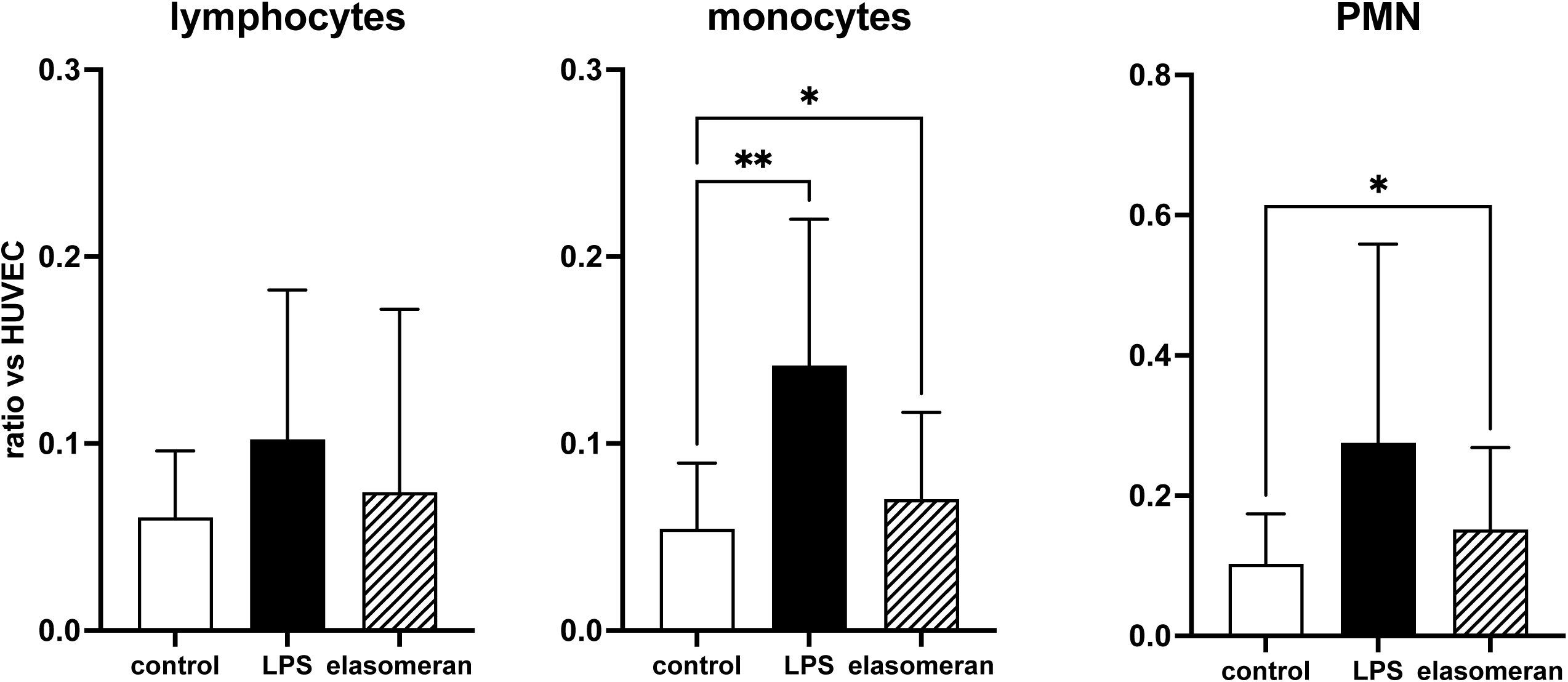
Flow cytofluorimetric evaluation of adhesion of leukocytes to HUVEC monolayers. White columns: negative control. Black columns: positive control (LPS 1µg/ml). Hatched columns: COVID-19 RNA vaccine Spikevax^TM^ (1µg/ml). Data are means±SD of 6-12 experiments. * = P<0.05 and ** = P<0.01 vs control.

### No evidence for the Spike protein gene sequence in the DNA isolated from HUVEC after incubation with the COVID-19 RNA vaccine SpikevaxTM

Real-time PCR analysis did not show any evidence for the presence of the Spike protein gene sequence in the DNA obtained from HUVEC exposed to 1 o 3 µg/ml of the COVID-19 RNA vaccine Spikevax^TM^, up to 144 h. As expected, a detectable signal was found when 5 µg/ml of cDNA, obtained from the reverse transcription of the COVID-19 RNA vaccine Spikevax^TM^, was added to HUVEC at the end of the incubation period (**Supplementary Table 3**).

## Discussion

The main results of the present study show how exposure of HUVEC to the COVID-19 RNA vaccine Spikevax^TM^ results in effective cell transfection, and subsequently endothelial cells produce consistent amounts of the Spike protein, which is secreted in the culture medium. Alongside with Spike protein production, HUVEC develop a proinflammatory phenotype, which includes increased gene expression of the proinflammatory cytokines IL-6 and TNF-α and of the adhesion molecules ICAM-1 and VCAM-1, resulting in increased attachment of leukocytes to HUVEC monolayers. The functional responses induced by the COVID-19 RNA vaccine Spikevax^TM^ in HUVEC in our study closely resemble the effects induced by exposure of HUVEC to the Spike protein of SARS-CoV-2 (Robles et al., 2022). In those experiments, incubation of HUVEC with whole Spike protein, its receptor-binding domain, or the integrin-binding tripeptide RGD induced the nuclear translocation of NF-κB and subsequent expression of the adhesion molecules VCAM-1 and ICAM-1, the coagulation factors TF and FVIII, the proinflammatory cytokines TNF-α, IL-1β, and IL-6, and ACE2, as well as the adhesion of leukocytes to the HUVEC monolayer, through mechanisms involving integrin *a*5β1 signaling (Robles et al., 2022).

In our model, the sequence of events, from HUVEC transfection to the production of Spike protein, to the stimulation of gene expression of proinflammatory cytokines and adhesion molecules, is quite rapid and tight, and occurs within the first hours of exposure to the vaccine. Peak expression of the genes for VCAM-1 and ICAM-1 occurs at 24 h, while for IL-6 and TNF-α it occurs at 48 h. Remarkably, Spike protein concentration in the culture medium at 24-48 h is in the order of 1.000-1.500 pg/ml. For comparison, the Spike protein EC_50_ reported by Robles et al. (2022) is about 300 ng/ml, which is 200-300 times higher than Spike protein concentrations in our model. A likely explanation for this apparent discrepancy is that, in our experiments, HUVEC themselves are the source of Spike protein, thus its concentration in the cell biophase, close to the HUVEC membrane, is many times higher than that measured after dilution in the entire culture medium. We further discuss later this fundamental concept.

HUVEC transfection is apparently transient, as suggested be the decline of the Spike protein RNA intracellular levels at 144 h in the presence of 1 µg/ml Spikevax^TM^. Nevertheless, with 3 µg/ml Spikevax^TM^, RNA levels were still at plateau. Based on our results, it is therefore not possible to say with certainty how long it takes for all RNA to be completely cleared from cells in this model.

Under the experimental conditions used in the present study, no evidence was obtained for the eventual integration of the Spikevax^TM^ gene sequence into the genome of HUVEC. Indeed, it has been shown that the sequence features of COVID-19 RNA vaccines seem to meet all the requirements for retroposition using L1 elements, the most abundant autonomously active retrotransposons in the human genome, a situation which makes the integration of RNA vaccines in the genomes of human cells theoretically possible (Domazet-Lošo, 2022). Remarkably, intracellular reverse transcription of the BioNTech–Pfizer COVID-19 RNA vaccine has been shown *in vitro* in the human liver cell line Huh7 (Aldén et al., 2022). Nevertheless, our experiments were performed on HUVEC at confluence, when most of the cells are quiescent G0, therefore in less favourable conditions to observe retroposition and integration. Different experimental conditions should be selected to properly test the potential genomic integration of the COVID-19 RNA vaccine Spikevax^TM^, for example in senescent HUVEC, where the expression of L1 sequences are significantly increased (Ramini et al., 2022).

Considering the present results, a crucial question is whether and to what extent these results can be translated to the clinical setting. In our experiments, we used the RNA of the COVID-19 vaccine Spikevax^TM^ at the concentrations of 1 and 3 µg/ml, which is a concentration range similar to that chosen by Schreckenberg et al. (2024) to study *in vitro* the direct effects of COVID-19 RNA vaccines on human cardiomyocytes, based on the biodistribution of [3H]-labelled lipid nanoparticles (LNP) RNA in rats (Schreckenberg et al., 2024), or by Cao et al. (2025) to study the effects of COVID-19 RNA vaccines on human macrophages (Cao et al., 2025). There are indeed no clinical studies describing the pharmacokinetics and the biodistribution of Spikevax^TM^ (or of any other COVID-19 RNA vaccines) in human subjects. Recently, however, the biodistribution of Spikevax^TM^ has been characterized in rats (Goody et al., 2026), where administration of 78 μg of RNA i.m. (260 μg/kg, about 153-fold the dose administered to a 60-kg human subject) resulted in an area under the curve (AUC) of 0,84-1,42 h x μg/ml, with estimated RNA C_max_ values in tissues with highest exposure, like draining lymph nodes and spleen, of 0,37 and 1,45 μg/g of tissue, respectively. Remarkably, such RNA levels resulted, on average, in 538 pg/ml of unbound SARS-CoV-2 Spike protein in plasma (Goody et al., 2026). This value can be compared to the 68±21 pg/ml of S1 Spike protein subunit found in the plasma of 11 out of 13 subjects receiving the COVID-19 RNA vaccine Spikevax^TM^, with peak values up to 150 pg/ml (Ogata et al., 2022), or to the 33.9±22.4 pg/ml of the free full-length Spike protein found in the plasma of a group of 16 adolescents who developed myocarditis after COVID-19 RNA vaccination, with peak values higher than 100 pg/ml (Yonker et al., 2023), or also to the 10,4 ng/ml (10.400 pg/ml) of circulating Spike protein found in a woman suffering from thrombocytopenia developed after vaccination with the COVID-19 RNA vaccine Spikevax^TM^ (Appelbaum et al., 2022). Although these are the only informations available on the plasma levels of the Spikevax^TM^-induced Spike protein in vaccinated subjects, values are superimposable to those found in rodents despite the higher RNA dose administered in the animal models (Goody et al., 2026). Most importantly, in our experiments in HUVEC cells exposed to 1 or 3 µg/ml of Spikevax^TM^ RNA, similar values, ranging between a few hundred and nearly 1.000 pg/ml of Spike protein are secreted in the cell culture medium in the first 24 h, when the effects on gene expression of the proinflammatory cytokines IL-6 and TNF-α and of the adhesion molecules ICAM-1 and VCAM-1 are already evident, suggesting the *in vivo* and clinical relevance of the results obtained in this *in vitro* model. In other terms, similar and even higher concentrations, in comparison to those found in our experiments, can occur in blood and, possibly, in tissues of human subjects receiving the standard dose of the COVID-19 RNA vaccine Spikevax^TM^ (Ogata et al., 2022; Yonker et al., 2023; Appelbaum et al., 2022).

It must be taken in mind that plasma levels of the Spike protein induced by the COVID-19 RNA vaccines likely result from leakage occurring in tissues where RNA-containing LNPs have localized and are therefore the site of protein production. According to the study by Goody et al. (2026), tissues such as injection site, draining lymph nodes, and spleen are highly exposed to the vaccine RNA, however virtually all tissues and organs are exposed, likely through the systemic circulation, including heart, liver, lung and even the brain. When evaluating the kinetics of vaccine-derived Spike protein in the body, it’s of key importance to take in mind that the protein originates from endogenous production, and its concentration is therefore likely higher in tissues where production occurs. For example, it is well established that levels of the neurotransmitter dopamine are up to 100 million times higher in brain areas where it is produced, in comparison to plasma where it occurs as a result of tissue spillover (Matt and Gaskill, 2020). It is reasonable to assume that the same occurs for COVID-19 vaccine-induced Spike protein, eventually leading to potentially toxic concentrations in tissues and organs where the Spike protein happens to be produced. Remarkably, at least a case has been reported describing a 76-year-old man with Parkinson’s disease who died three weeks after receiving his third COVID-19 vaccination (first dose with ChAdOx1 nCov-19 vector vaccine, second and third doses of the BNT162b2 RNA). The autopsy detected the Spike protein within the foci of inflammation in both the brain and the heart, particularly in the endothelial cells of small blood vessels, and, since no SARS-CoV-2 nucleocapsid protein could be detected, the presence of the Spike protein has been ascribed to vaccination rather than to viral infection (Mörz, 2022).

The virtually random localization of RNA/LNP complexes in different organs and tissues is possibly one of the major determinants of COVID-19 RNA vaccines-induced toxic effects. For example, the excess amount of free Spike protein in postvaccine myocarditis could well result from excessive production because of factors specific to the recipient organism, such as more efficient protein synthesis, especially in people of younger ages, or localization of COVID-19 vaccine RNA/LNP complexes in tissues or organs with intrinsically high protein synthesis capacity (e.g., liver, ileum, heart, skeletal muscle) (Anisimova et al., 2020; Knudsen and Prasad, 2023; Cosentino and Marino, 2023), resulting in excessive protein production for too long or in tissues or organs sensitive to its toxicity, in any case overcoming the binding or buffer capacity of the vaccine-induced antibody response (Cosentino and Marino, 2022).

An excessive and dystopian production of the Spike protein induced by the COVID-19 vaccine could effectively explain the studies documenting vaccine-induced adverse effects in relation to plasma Spike protein levels. For example, in the above-mentioned case of the woman with Spikevax^TM^– induced thrombocytopenia, the very high vaccine-induced Spike levels in plasma 10 days after vaccination suggests a high protein synthesis capacity of the tissues where the RNA/LNP complexes have located and/or a localization close to the systemic circulation, resulting in high quantities of Spike protein released into the blood, which in turn directly affected circulating platelets (Appelbaum et al., 2022). On the other side, the levels reported in the study by Yonker et al. (2023) in association with the onset of myocarditis could reasonably indicate that the primary site of production of the Spike protein is precisely the cardiac parenchyma, thus exposing the myocardial tissue to higher and therefore toxic concentrations of Spike protein. Indirect support to this possibility comes from studies showing the presence of the vaccine-induced Spike in endomyocardial biopsies from patients with post-COVID-19 vaccine myocarditis up to nearly 2 months after COVID-19 vaccination (Baumeier et al., 2022).

The present results, showing that a COVID-19 RNA vaccine can transfect human endothelial cells, inducing a proinflammatory response resulting in increased leukocyte adhesion, is in line with clinical evidence documenting inflammation and endothelial dysfunction following vaccination. Terentes-Printzios et al. (2022) described a prominent increase in blood levels of high-sensitivity C-reactive protein and a transient deterioration of endothelial function, assessed by brachial artery flow-mediated dilatation, in a cohort of otherwise healthy subjects 24 h after vaccination with the COVID-19 RNA vaccine BNT162b2. Impaired flow-mediated dilatation two weeks after the second dose of the COVID-19 RNA vaccine BNT162b2 was reported also in a group of medical staff members at Hiroshima University Hospital (Yamaji et al., 2024). Another study of particular interest (Castro-Robles et al., 2025) reports increased serum levels of soluble VCAM-1 and endocan following the third dose of the COVID-19 RNA vaccine BNT162b2, possibly in association with reduced Spike protein-specific antibody production. Soluble VCAM-1, shedding from the surface of endothelial cells during inflammatory responses, is a biomarker of immunological diseases, cancer, autoimmune myocarditis, and a predictor of mortality and morbidity in patients with chronic heart failure, endothelial injury in patients with coronary artery disease, and arrhythmias (Troncoso et al., 2021), while endocan, also known as endothelial cell specific molecule-1, is a soluble proteoglycan secreted by endothelial cells, and elevated plasma levels reflect endothelial activation and dysfunction, that may be related to cardiovascular disease such as hypertension, diabetes, angina pectoris and acute myocardial infarction (Chen et al., 2022). Our results provide a mechanistic explanation to such observational reports, supporting the notion that COVID-19 RNA vaccines directly affect endothelial cells, inducing a proinflammatory response which in turn may represent a risk factor for immune and inflammatory cardiovascular disease. While myocarditis and pericarditis are already included among the possible adverse effects of COVID-19 RNA vaccines (Buoninfante et al., 2024), a causal role of these vaccines in pathologies resulting from inflammatory endothelial dysfunction, for example during atherosclerosis, autoimmune inflammation, and systemic inflammatory conditions, should be taken into consideration.

The major limitation of our study lies in the limited availability of vaccine for *in vitro* experiments. The product used in our experiments was a leftover material from the Italian National Institute of Health (ISS). However, COVID-19 RNA vaccines are formally licensed for marketing in Italy but are not available in pharmacies. Therefore, it is extremely difficult to obtain products for *in vitro* and *in vivo* pharmacological and functional studies. For this reason, in our experiments we had to limit ourselves to using two concentrations, and we were unable to construct complete concentration-response curves or to test the vaccine’s effect in the presence of other pharmacological agents. Consequently, for example we can only guess that the effects that we observed in HUVEC involve integrin *a*5β1 signaling, based on published studies which showed that the effects of the SARS-CoV-2 Spike protein on HUVEC is sensitive to inhibitors of α5β1, such volociximab and ATN-161 (Robles et al., 2022). In addition, we were unable to test in HUVEC the possible effects of agents which have been shown to counteract the detrimental effects of the SARS-CoV-2 Spike protein, such as nattokinase (Tanikawa et al., 2022), aspirin (Ciszewski et al., 2024), or genistein (Cao et al., 2025). Further limitations include the failure to distinguish between the possible effects of the different vaccine components. Although the literature suggests that the proinflammatory effect of RNA vaccines should be primarily ascribed to the protein, it is important to remember that modified RNAs also have specific properties (Rubio-Casillas et al., 2024). Furthermore, the ability of LNPs to induce neutrophil activation and neutrophil extracellular trap (NET) formation, resulting in the establishment of a pro-metastatic niche, has recently been described in a rodent model (Wang et al., 2026).

In conclusion, the present study shows that the COVID-19 RNA vaccine Spikevax^TM^ can transfect human endothelial cells, inducing a proinflammatory response and resulting in increased leukocyte adhesion. Results provide a mechanistic explanation to post-COVID-19 RNA vaccination pathologies resulting from endothelial dysfunction.

## Acknowledgements

The authors are grateful to Ilaria Muller, Fondazione IRRCS Ca’ Granda Ospedale Maggiore Policinico, Department of Endocrinology, Milan, Italy, and to Chiara Di Meo, Department of Drug Chemistry and Technologies, Sapienza University of Rome, 00185 Roma, Italy, for the insightful discussions during the design and execution of the experiments, and the interpretation of the results of this study.

## Availability of data and materials

The datasets used and/or analyzed during the current study are available from the corresponding author on reasonable request.

## Authors’ contributions

MC, FM, and GVF contributed to the study conception and design. ER and AL were responsible for HUVEC experiments, MF, NS, and ML performed molecular biology assays. All authors contributed to the analysis and interpretation of data and were involved in drafting the article or revising it critically for important intellectual content. All authors approved the final version to be published and agree to be accountable for all aspects of the work in ensuring that questions related to the accuracy or integrity of any part of the work are appropriately investigated and resolved and declare to have confidence in the integrity of the contributions of their co-authors.

