## Supplementary Tables for "The COVID-19 RNA vaccine Spikevax^TM^ transfects human umbilical endothelial cells and induces Spike protein production, inflammation and leukocyte adhesion"

**Supplementary Table 1.** Real-Time PCR Primers for SARS-CoV-2 T expression.

| **Gene Symbol** | **UniGene ID** | **Interrogated Sequence** *RefSeq/GenBank mRNA* | **Detected Coding Transcripts** | **Amplicon Context Sequence** | **Chromosome Location** | **Amplicon Lenght** | **Annealing temperature (°C)** | **Efficiency** (%) |
|---|---|---|---|---|---|---|---|---|
| **SARS-CoV-2 S gene** | Vi07918636_s1 | Patented by Thermo Fisher Scientific (Waltham, MA, USA) | | | | | 60 | 99 |
| **TNF** | Hs.241570 | NC_000006.11<br>NG_007462.1<br>NG_012010.1<br>NT_007592.15<br>NT_113891.2<br>NT_167244.1<br>NT_167245.1<br>NT_167246.1<br>NT_167247.1<br>NT_167248.1<br>NT_167249.1 | ENST00000328965<br>ENST00000445232<br>ENST00000594551<br>ENST00000443707<br>ENST00000412275<br>ENST00000449264<br>ENST00000577810<br>ENST00000326294 | GGGGTCTTCCAG<br>CTGGAGAAGGGT<br>GACCGACTCAGC<br>GCTGAGATCAAT<br>CGGCCCGACTAT<br>CTCGACTTTGCC<br>GAGTCTGGGCAG<br>GTCTACTTTGGG<br>ATCATTGCCCT<br>GTGAGGAGGACG<br>AACATC | 6:31545204-31545328 | 95 | 60 | 99 |

| | | | |
|---|---|---|---|
| | | | ENST00000448781<br>ENST00000420425<br>ENST00000394126<br>ENST00000356271<br>ENST00000394128<br>ENST00000394127<br>ENST00000422942<br>ENST00000501516<br>ENST00000536318<br>ENST00000431269<br>ENST00000376122<br>ENST00000383496<br>ENST00000264203<br>ENST00000375144<br>ENST00000375142<br>ENST00000401084<br>ENST00000439554 |

| | | | | | | | | |
|---|---|---|---|---|---|---|---|---|
| **IL-6** | Hs.654458 | NC_000007.13 NG_011640.1 NT_007819.17 | ENST00000404625 ENST00000426291 ENST00000401651 ENST00000407492 ENST00000401630 ENST00000406575 ENST00000258743 ENST00000420258 | GTATACCTAGAGTACCTCCAGAACAGATTTGAGAGTAGTGAGGAACAAGCCAGACTGTGCAGATGAGTACAAAAGTCCTGATCCAGTTCCTGCAGAAA | 7:22769178-22769276 | 69 | 60 | 98 |
| **ICAM1** | Hs.643447 | NC_000019.9 NG_007728.1 NG_012083.1 NT_011295.11 | ENST00000264832 ENST00000423829 | GCATTGTCCTCAGTCAGATACAACAGCATTTGGGGCCATGGTACCTGCACACCTAAAACACTAGGCCACGCATCTGATCTGTAGTCACATGACTAAGCCAAGAGGAAGGAGCAAGACTCAAGACATGA | 19:10396090-10396217 | 98 | 60 | 99 |
| **VCAM1** | Hs.109225 | NC_000001.10 NG_023034.2 NT_032977.9 | ENST00000370119 ENST00000347652 | GGAATTAACCAGGCTGGAAGAAGCAGAAAGGAAGT | 1:101198189-101200172 | 137 | 60 | 102 |

| | | | | | | | | |
|---|---|---|---|---|---|---|---|---|
| | | | ENST00000294728<br>ENST00000370115 | GGAATTAATTATC<br>CAAGTTACTCCA<br>AAAGACATAAAA<br>CTTACAGCTTTTC<br>CTTCTGAGAGTG<br>TCAAAGAAGGAG<br>ACACTGTCATCAT<br>CTCTTGTACATGT<br>GGAAATGTTCCA<br>GAAACATGGATA<br>ATCCTGAA | | | | |
| **RPS18** | Hs.627414 | NC_000006.11<br>NT_007592.15<br>NT_113891.2<br>NT_167245.1<br>NT_167247.1<br>NT_167248.1<br>NT_167249.1 | ENST00000454021<br>ENST00000486781<br>ENST00000484321<br>ENST00000211372<br>ENST00000477055<br>ENST00000476288<br>ENST00000439602<br>ENST00000474973<br>ENST00000457341<br>ENST00000494232<br>ENST00000434122 | GTGGAACGTGTG<br>ATCACCATTATGC<br>AGAATCCACGCC<br>AGTACAAGATCC<br>CAGACTGGTTCT<br>TGAACAGACAGA<br>AGGATGTAAAGG<br>ATGGAAAATACA | 6:33243742-<br>33243838 | 67 | 60 | 98 |

**Supplementary Table 2.** Flow cytometry analysis of HUVEC viability after incubation for 144 h with the COVID-19 RNA vaccine elasomeran. Anx: annexin; PI: propidium iodide. Data are percentages of total cells in the cell sample and are expressed as means±SD of n = 3 independent replicates.

| **Cells** | **control** | **elasomeran** 1 µg/ml | **elasomeran** 3 µg/ml |
|---|---|---|---|
| viable (Anx-PI-) | 90,1±2,1 | 91,2±3,0 | 92,1±1,0 |
| early apoptotic (Anx+PI-) | 5,6±1,4 | 4,9±1,8 | 3,8±0,5 |
| late apoptotic (Anx+PI+) | 4,2±0,7 | 3,7±1,3 | 3,8±0,5 |
| necrotic (Anx-PI+) | 0,1±0,1 | 0,1±0,0 | 0,2±0,1 |

**Supplementary Table 3.** Real-time PCR assay of the Spike protein gene sequence in the DNA isolated from HUVEC after incubation with the COVID-19 RNA vaccine elasomeran. NC, negative control; HKG, housekeeping gene; nd: not detected. Data are cycle thresholds and are expressed as means±SD of n = 2 independent replicates.

| **incubation time** (h) | **elasomeran** 1 µg/ml | **HKG** | **NC** | **HKG** | **elasomeran** 3 µg/ml | **HKG** | **NC** | **HKG** | **elasomeran cDNA** 5 µg/ml | **HKG** |
|---|---|---|---|---|---|---|---|---|---|---|
| 2 | nd | 13.1±0.2 | nd | 13.2±0.2 | nd | 11.2±0.3 | nd | 12.4±0.1 | 13.2±0.5 | 30.9±0.2 |
| 4 | nd | 13.5±0.2 | nd | 13.4±0.4 | nd | 11.3±0.1 | nd | 12.1±0.0 | 12.9±0.1 | 31.2±0.6 |
| 24 | nd | 13.2±0.2 | nd | 13.0±0.0 | nd | 11.2±0.3 | nd | 12.2±0.1 | 12.8±0.4 | 31.2±0.2 |
| 48 | nd | 13.5±0.3 | nd | 13.3±0.1 | nd | 11.3±0.5 | nd | 12.1±0.1 | 13.2±0.2 | 30.7±0.4 |
| 72 | nd | 13.2±0.1 | nd | 13.3±0.1 | nd | 11.3±0.4 | nd | 12.1±0.1 | 12.7±0.0 | 31.2±0.3 |
| 144 | nd | 13.2±0.2 | nd | 13.3±0.1 | nd | 11.3±0.2 | nd | 12.1±0.1 | 13.1±0.4 | 30.8±0.1 |
