## Supplementary Figures for "The COVID-19 RNA vaccine Spikevax^TM^ transfects human umbilical endothelial cells and induces Spike protein production, inflammation and leukocyte adhesion"

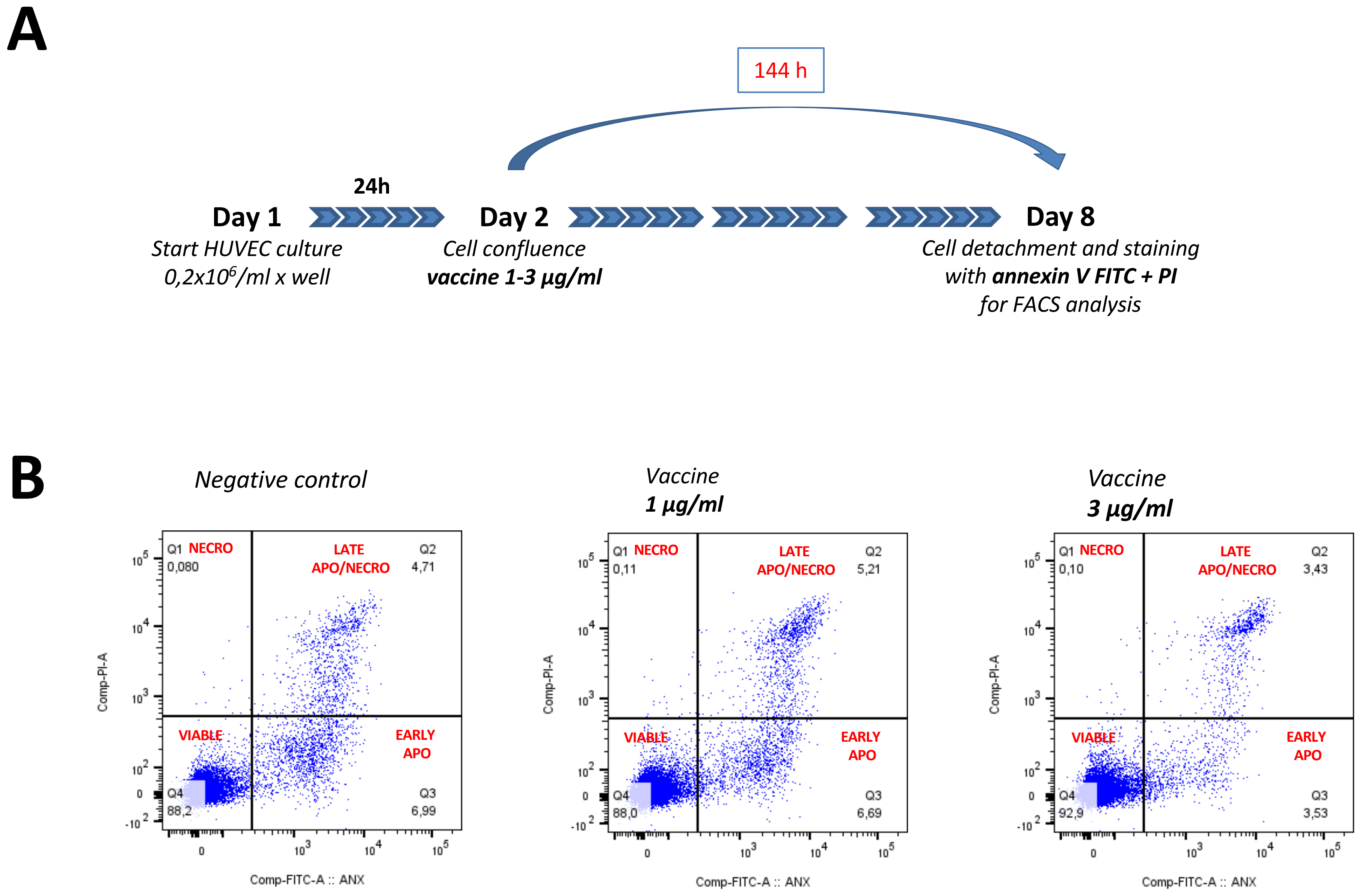


**Supplementary Figure 1.** Flow cytometry analysis of HUVEC viability. Panel A: experimental design. Panel B: results from a representative experiment.

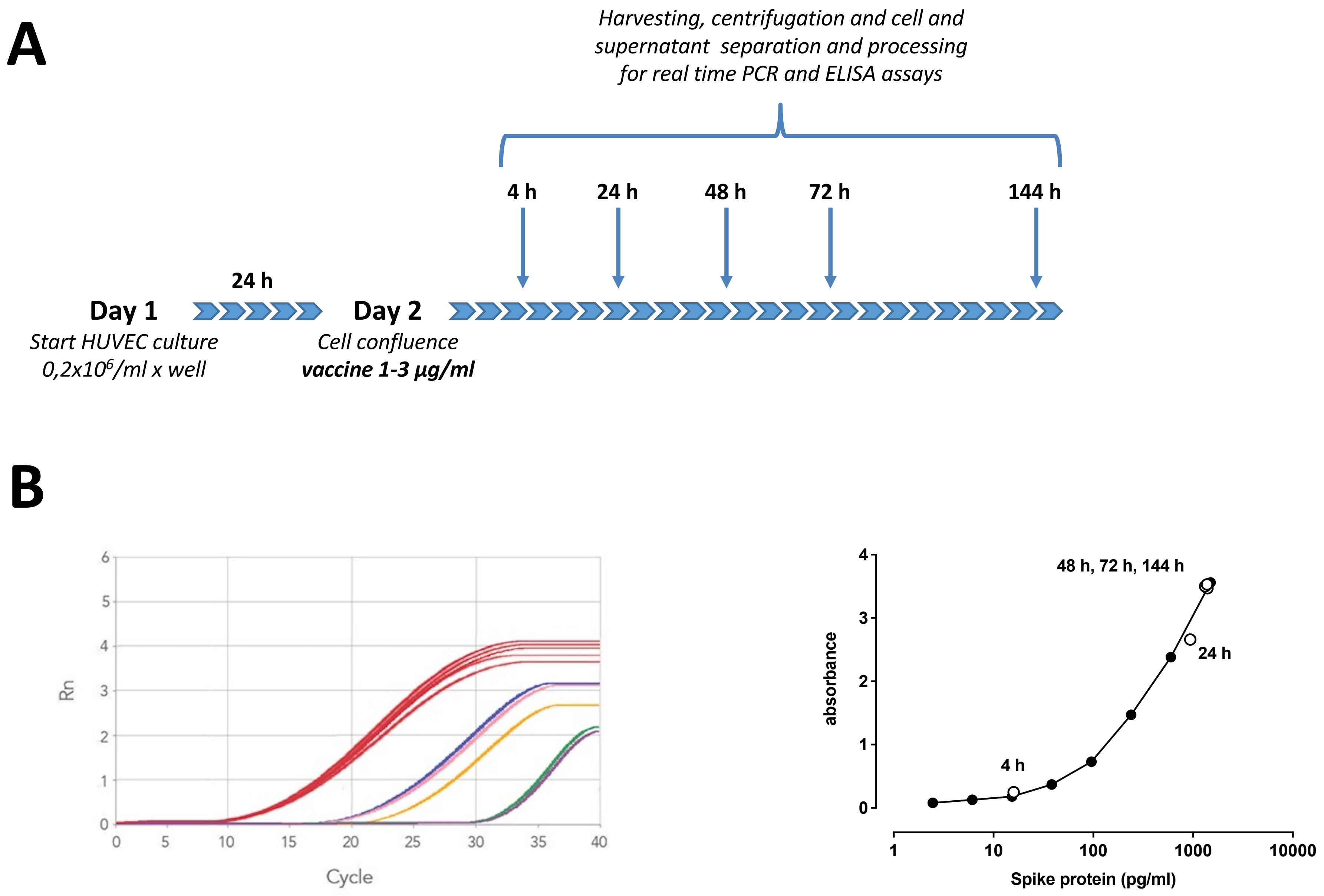


**Supplementary Figure 2.** Spike protein expression in HUVEC cultured in the presence of vaccine at different concentrations. Panel A: experimental design. Panel B: results from a representative experiment with 1 µg/ml vaccine. Left: PCR assay; red, 18sRNA (housekeeping gene), green, 4 h; yellow, 24 h; blue, 48 h; pink, 72 h; purple, 144 h. Right: ELISA assay; black circles, calibration curve; white circles: experimental samples.

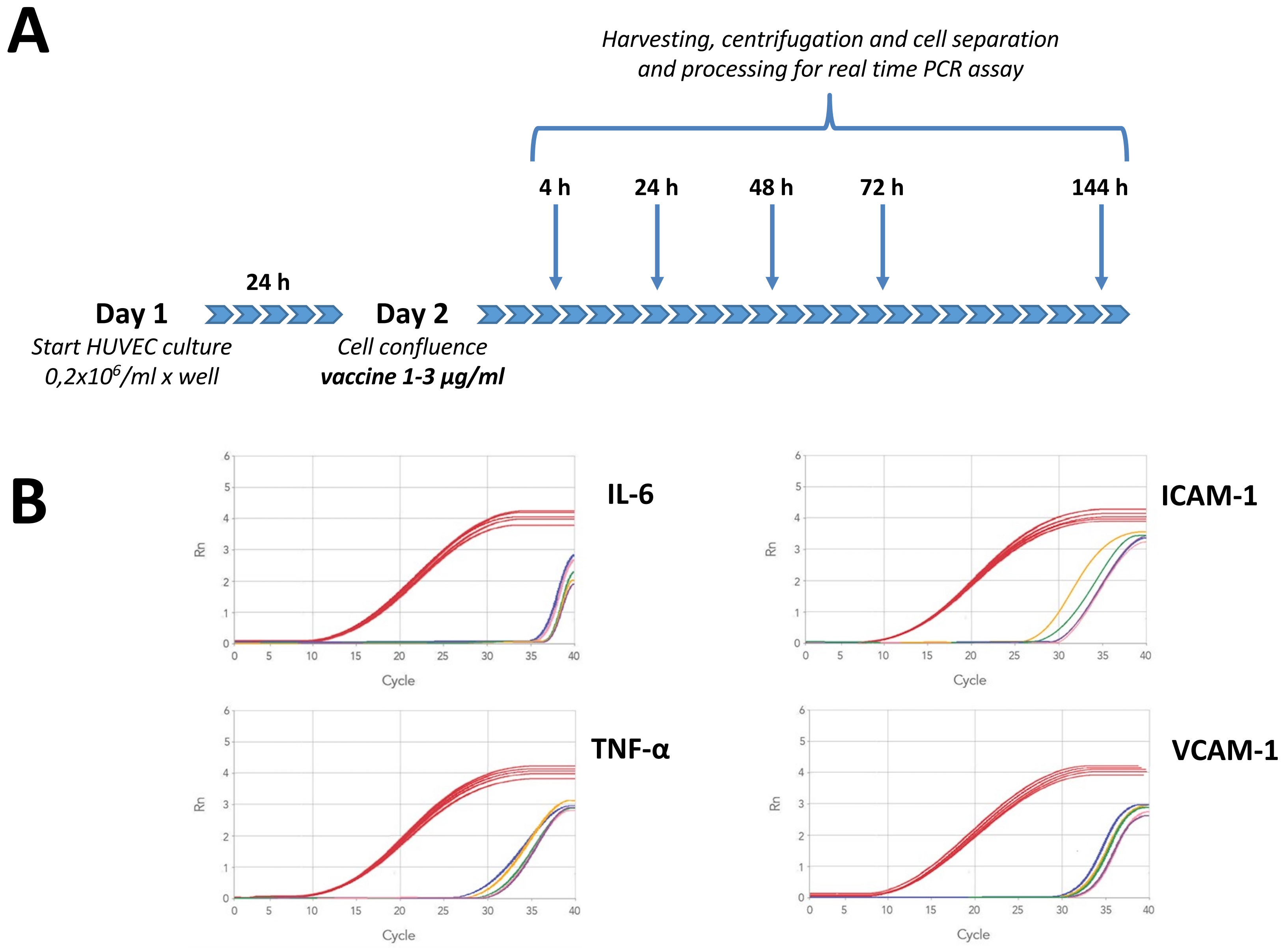


**Supplementary Figure 3.** IL-6, TNF-α, ICAM-1 and VCAM-1 mRNA levels in HUVEC cultured in the presence of vaccine at different concentrations. Panel A: experimental design. Panel B: results from a representative experiment with 1 µg/ml vaccine. Left: PCR assay; red, 18sRNA (housekeeping gene), green, 4 h; yellow, 24 h; blue, 48 h; pink, 72 h; purple, 144 h. Right: ELISA assay.

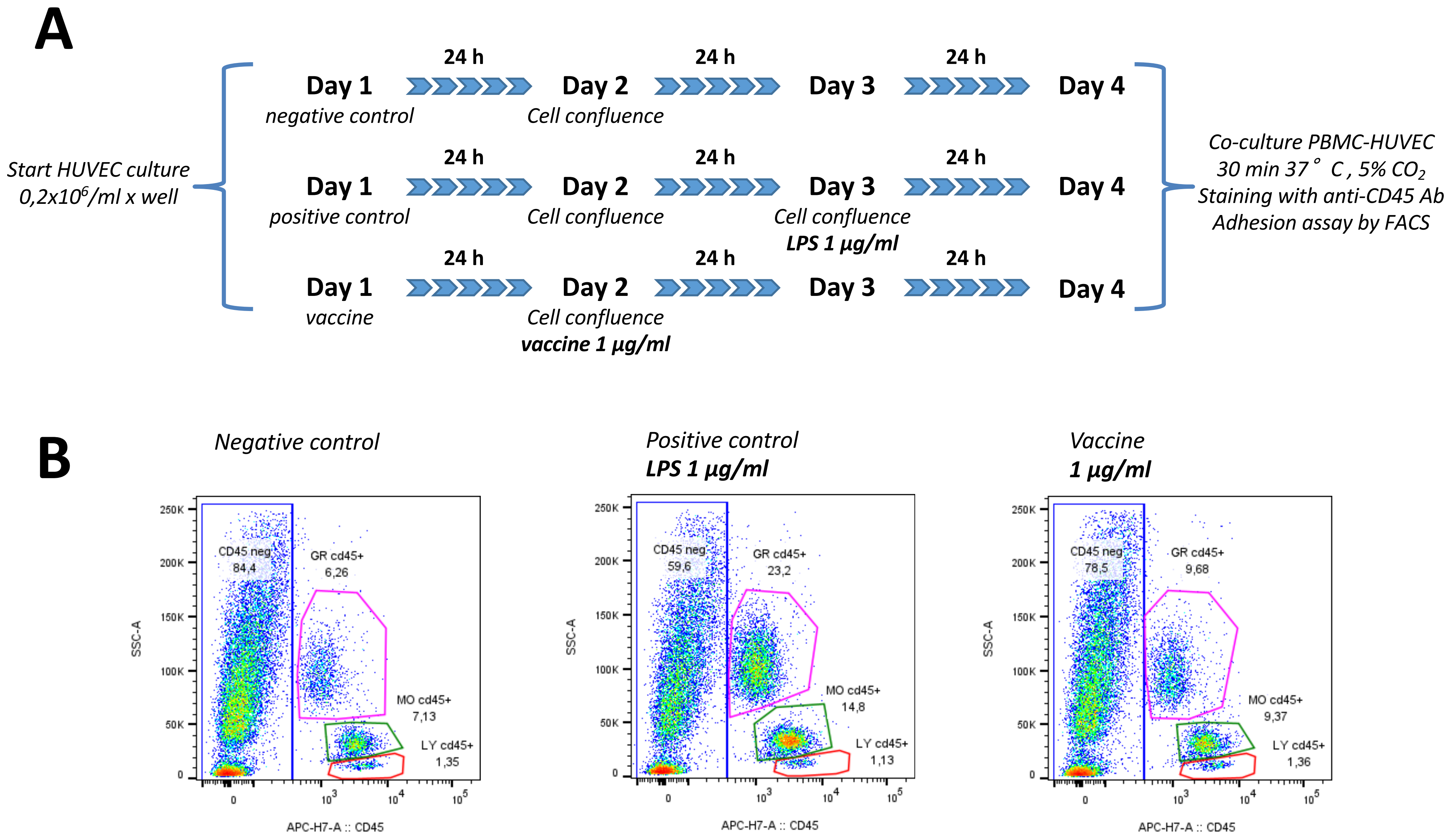


**Supplementary Figure 4.** Effect of vaccine on the adhesion of human leukocytes to HUVEC monolayers, assessed by flow cytometry. Panel A: experimental design. Panel B: results from a representative experiment.

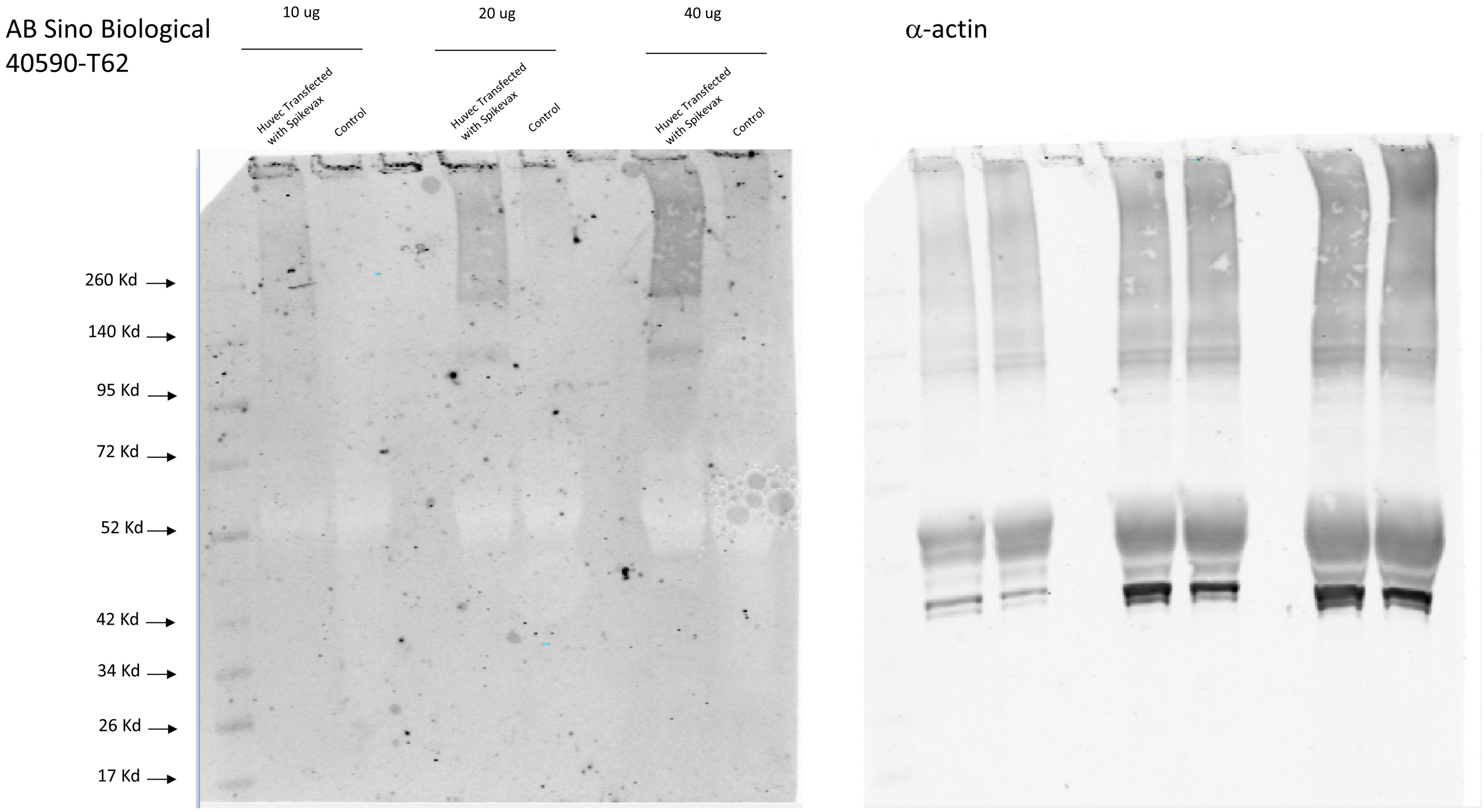


**Supplementary Figure 5.** Complete Western blot of total protein extracts from HUVEC treated for 48 h with the COVID-19 RNA vaccine Spikevax™ at the concentration of 3 µg/ml. Control: untreated HUVEC.
